# Iterative remodeling of the mouse uterus requires Hedgehog signaling

**DOI:** 10.1101/2022.10.27.514107

**Authors:** Elle C. Roberson, Ngan Kim Tran, Anushka N. Godambe, Trinity Rust, Michelle Nguimtsop, Harrison Mark, Rebecca D. Fitch, John B. Wallingford

## Abstract

The adult uterus regenerates in the human during the menstrual cycle, and remodels in the mouse during the estrous cycle. Decades of work has demonstrated that this process is controlled by cycling steroid hormones, estrogen and progesterone. However, downstream signaling pathways that link hormonal action to this regeneration and remodeling are yet to be identified in the cycling uterus. We set out to identify these pathways, with the overarching hypothesis that developmental signaling pathways are redeployed in the adult uterus to control remodeling in the mouse. We were surprised to find that the majority of Hedgehog signaling components were transcriptionally co-regulated throughout the estrous cycle. To test the role of Hh signaling in cyclical uterine remodeling, we conditionally knocked out the major activator of the pathway, *smoothened* (*Smo*) using the *progesterone receptor cre* (*PR-Cre*). In the absence of Hh signaling, the uterus no longer remodels throughout the estrous cycle. We also show that the smooth muscle fibers of the uterus are significantly larger in the conditional knockouts compared to the controls suggesting hypertrophy of the smooth muscle. Our findings support the possibility that this smooth muscle homeostasis may underlie important aspects of uterine function such as contractility during late-stage pregnancy or the development of uterine smooth muscle tumors.

## Introduction

Unlike other adult tissues, the uterus repeatedly undergoes extensive remodeling during the menstrual and estrous cycle, implantation, and pregnancy. While both uterine endometrium (composed of epithelium and fibroblasts) and myometrium (the outer layers of smooth muscle) remodel during pregnancy and parturition, only the endometrium remodels during the menstrual and estrous cycle ^1^. Decades of research has described the role of hormones – like estrogen and progesterone – in the remodeling of the adult uterus. For example, in animals that menstruate (∼1.6% of eutherian mammals), the endometrium proliferates during the first half of the cycle – the proliferative phase, when estrogen increases, and then differentiates in the second half of the cycle – the secretory phase, when progesterone increases ^2^. This cycle of proliferation followed by differentiation prepares the uterus for implantation of a fertilized embryo ^2^. If implantation does not occur, then the endometrium is shed during menstruation ^2^. By contrast, in animals that don’t menstruate (98.4% of eutherian mammals), the endometrium thickens during the first half of the estrous cycle – stages proestrus and estrus, when estrogen increases, and then dramatically thins in the second half – stages metestrus and diestrus, when progesterone increases ^3,4^. While the morphological changes have been well characterized, the downstream signaling pathways that control cyclical uterine remodeling have received little attention.

In contrast to cyclical uterine remodeling, the signaling required for implantation in mouse models has been well investigated. For example, a direct target of progesterone signaling during implantation is *Indian hedgehog* (*Ihh*) ^5-8^. At implantation, Ihh is secreted by the luminal epithelium and drives stromal cell proliferation and differentiation ^8-10^. When *Ihh* is conditionally knocked out in the adult uterus, female mice are infertile due to implantation failure ^8^. In contrast, when a constitutively active *Smoothened, SmoM2*, is expressed in the uterus, implantation also fails ^11^. In addition to these mouse models, several single cell, transcriptomic, and cistromic datasets have described the downstream molecular genetic control of implantation in the mouse ^5,12-16^.

Transcriptomic and single cell datasets of the cycling uterus in humans have likewise illuminated the molecular control of regeneration during the menstrual cycle ^17-21^. However, we lack a comprehensive understanding of the molecular cascades that drive cyclical uterine remodeling in any genetically tractable model system. While some single cell and transcriptomics studies have been performed using the mouse ^22,23^, and others have reported the expression of individual genes or gene families, the absence of a dynamic mouse uterine transcriptome at each stage of the normal estrous cycle represents a critical gap in our knowledge.

The present study documents the transcriptomic landscape of the whole mouse uterus across the estrous cycle, revealing the patterned expression of a variety of signaling pathways more commonly associated with embryonic development, including Hedgehog (Hh) signaling. Here we investigate how Hh signaling regulates uterine remodeling during the estrous cycle. Similar to the studies at implantation, we show that Hh signaling is required for endometrial remodeling across the estrous cycle. We also show that Hh signaling maintains homeostasis of the myometrium, which suggests that Hh signaling may play a role in other aspects of myometrial biology, like pregnancy and parturition, or the progression of uterine smooth muscle tumors.

## Materials and Methods

### Mice

6-8-week-old Swiss Webster female mice were obtained from Charles River, while *progesterone receptor cre* (*PR-Cre*, Stock: 017915) ^24^ and *smoothened* floxed (*Smo*^*fl*/+^, Stock: 004526) ^25^ animals were obtained from Jackson Labs. Mice were housed in individually ventilated cages in a pathogen-free facility with continuous food and water, with a controlled light cycle (light from 7am-7pm). Floxed females were bred to *PR-Cre*^*+/-*^ males to produce experimental (*PR-Cre*^+/-^; *Smo*^*fl*/*fl*^, cKO) or control females. 7-12-week-old females were estrous cycle staged using standard vaginal cytology ^26^. Mice were humanely euthanized with extended CO_2_ exposure followed by cervical dislocation, and female reproductive tracts were dissected. All animal experiments were approved by the University of Texas at Austin Institutional Animal Care and Use Committee.

### Tissue processing, histology, and immunofluorescence

Dissected uteri were gently affixed to a strip of index card to keep the tissue straight, and fixed in 4% paraformaldehyde for either 4-6hr at room temperature (RT) or overnight at 4°C. Fixed uteri were washed in PBS, placed in 70% ethanol (EtOH) for at least 24hr, paraffin embedded, and sectioned to produce 5μm sections. Sections were allowed to dry overnight.

For Hematoxylin and Eosin (H&E) staining, slides were baked at 60°C for 20min. Slides were de-waxed with three 5min xylene incubations, then rehydrated (100% EtOH, 3min x 2; 95% EtOH, 2min; and ddH2O, 4min). After rehydrating, slides were treated as follows: incubated in hematoxylin (3min), running tap water (1min), differentiated (45sec), water (30sec), bluing reagent (1min), water (30sec), 80% EtOH (1min), Eosin Y (3min), dehydrated (95% EtOH, 1min x 2; 100% EtOH, 3min x 2; xylene, 3min x 2), coverslipped with Cytoseal and allowed to cure overnight. H&E stained sections were imaged using a Keyence microscope (BZ-X 710).

For Masson Trichrome staining, we used a Statlab kit (catalog #: KTMTRLT) and followed their instructions with some exceptions. In brief, slides were baked at 60°C for 20min, de-waxed in xylene (2×5min), and rinsed in 100% ethanol (3×1min) followed by running tap water (1min). Slides were then incubated in Bouin’s Fluid (RT, overnight), rinsed in running tap water (3min), immersed in Hematoxylin (5min), rinsed in running tap water (2min), immersed in Biebrich Scarlet-Acid Fusion (1min), rinsed in running tap water (45min), immersed in Phosphomolybdic/ Phosphotungstic Acid (15min), then Aniline Blue stain (10min), rinsed in running tap water (1min), and then immersed in 1% Acetic Acid (5min). Finally, slides were dehydrated in 100% ethanol (3×1min), cleared with xylene (3×1min), and mounted with Cytoseal. Masson Trichrome stained slides were also imaged with the Keyence microscope.

### Image quantitation

At least four sections 100μm apart were imaged and quantified in order to accurately assess tissue morphology differences, and all quantitation was performed in FIJI. Keyence images were calibrated in FIJI based on the length of the scale bar. Uterine area was quantified in an unbiased manner by thresholding for uterine tissue and measuring the thresholded area. The lumen area was calculated by tracing along the apical edge of lumen epithelium. The stromal area was calculated by tracing along the stromal-myometrium junction and subtracting the lumen area. Uterine glands were counted with the multipoint tool, as previously described^4^. Longitudinal smooth muscle thickness was quantified by drawing a straight line from the outer edge of the circular muscle to the outer edge of the longitudinal muscle. Muscle fiber size and number was quantified using only Masson Trichrome staining, and the analysis started in Ilastik using the Pixel Classification mode ^27^. First, we trained Ilastik on small images from each of our samples, classifying pixels as either muscle, collagen, or not-applicable. After training, Ilastik analyzed whole images to specify pixels and produced masks of the classification. Using FIJI we were able to focus specifically on the muscle mask and quantify both number and size using the Analyze Particles function.

### RNA isolation and cDNA synthesis

For 3’ TagSeq (see below) of WT tissue, uterus samples were collected in triplicate at all estrous cycle stages. For qPCR, WT samples were collected (n=7) for each estrous cycle stage, and control and cKO samples (n=4) for estrus and diestrus. Following storage in RNA*Later* Storage Solution (Sigma, cat#: R0901) at -20°C, uterine tissue was physically lysed using a Beadbug 6 Microtube Homogenizer (Sigma-Aldrich, cat#: Z742682) and the lysate was spun through a QIAshredder column (Qiagen, cat#: 79656) to fully homogenize. A Qiagen RNeasy mini kit (Qiagen, cat#: 74106) was used to harvest RNA for RNAseq and qPCR. Total RNA was then either provided to the Genomic Sequencing and Analysis Facility at the University of Texas at Austin for 3’ TagSeq, or cDNA was synthesized using the iScript Reverse Transcription SuperMix (BioRad, cat#: 1708841) for qPCR.

### qPCR

Most primers were designed from a database for mouse and human qPCR primers incorporated into the UCSC genome browser ^28^. See Supplemental Table S-1 for primer details. We confirmed specificity of primers by ensuring that they BLAST to no more than 1 site in the genome and we only used primers whose melting curve displayed a single peak.

Primer sets were diluted from a stock (100μm in TE buffer) to 1μM in distilled deionized H_2_O. 2μL of each primer set (in technical quadruplicate) was allowed to dry in the bottom of a well in a MicroAmp Fast Optical 96 Well Reaction Plate (Thermofisher, cat#: 43-469-06). 10μL of a master mix of cDNA (250pg/well), SYBR Select Master Mix (Thermofisher, cat#: 44-729-18), and distilled deionized water was added to each well, the plates were sealed with MicroAmp Optical Adhesive Film (Thermofisher, cat#: 43-119-71) and allowed to incubate at RT in the dark for at least 15min to rehydrate primer. Plates were run on a ViiA-7 Real-Time PCR system (ThermoFisher), and CT values were auto-determined by the ViiA-7 software. The standard 2^-x0394;ΔCt^method was then used to determine fold change based on the geomean of three ‘housekeeping’ genes (*Hprt, Dolk*, and *Sra1*) ^29,30^.

### TagSeq

Tissue samples were collected in triplicate for each of the four estrous stages, accounting for 12 samples in total, and total RNA was collected as described above. Library preparation and sequencing for TagSeq ^31,32^, a form of 3’ RNA sequencing, were performed by the Genomic Sequencing and Analysis Facility (GSAF) at The University of Texas at Austin. Total RNA was isolated from each sample by addition of Trizol (Thermo Fisher) and the sample was transferred to a Phasemaker tube (Thermo Fisher). Total RNA was extracted following the protocol supplied by the manufacturer and further cleaned up using a RNeasy MinElute Cleanup Kit (Qiagen). RNA integrity number (RIN) was measured using an Agilent Bioanalyzer and 100ng of RNA was used for the TagSeq protocol. The fragmentation/RT mix was prepared and added to each RNA sample, then heated to 95°C for 2.5 minutes on a Thermocycler and immediately put on ice for 2 minutes. After cooling and addition of the template switching oligo and SmartScribe RT, the fragmented RNA reaction was incubated at 42°C for 1hr, 65°C for 15 min. Next an AmPure bead clean-up was completed for the cDNA before it was amplified to incorporate the Illumina sequencing primer site, followed by another cleanup. The remaining portions of the Illumina adapter (the i5 and i7 indices) were then added through an additional 4 cycles of PCR. Final libraries were quantified with PicoGreen then pooled equally for size selection using the Blue Pippin from 355-550 bp. Resulting libraries were sequenced using an Illumina HiSeq 2500 instrument (50-nt single reads).

### Sequence data pre-processing

Sequencing data quality was evaluated using the FastQC tool (v0.11.9) ^33^ and reports were aggregated with the MultiQC program (v1.0) ^34^.

### TagSeq data analysis

Single-end pseudo-alignment was performed against the mouse transcriptome (GENCODE M25 transcript sequences) using kallisto (v0.45.0) ^35^ with options -l 200 -s 50 --single-overhang –bias. Downstream analysis of transcript abundance data was performed in R following protocols outlined in Bioconductor ^36^. The tximport package ^37^ was used to roll up transcript-level counts into gene-level counts provided to the DESeq2 package ^38^. Count data matrices were filtered to remove genes with fewer than 1 read across all included samples. We used estrous stage as the model to explore gene expression across the cycle. Differentially expressed gene (DEGs) results reported are those with maximum adjusted p-value 0.05 and log2 fold change greater than 1.0 or less than -1.0. Heatmaps were made using z-scores based on the VSD and ComplexHeatmap ^39^ in R. Volcano plots were made according to the DESeq2 vignette.

Full DESeq2 results are provided in at the GEO accession number GSE216120.

## Results

### The murine uterus remodels across the estrous cycle

Previous reports have described changes to uterine and endometrial thickness and gland number during the mouse estrous cycle ^3,4^, but other parameters have not been reported. To gain deeper insights, we visualized the gross tissue morphology of uteri at each estrous cycle stage using H&E staining (**Fig. 1A-D**, n=8 per stage), and then quantified different uterine compartments including the whole uterus, lumen, endometrium, and myometrium. The uterus is thickest at proestrus and then steadily decreases in area until it becomes thinnest at diestrus (**Fig. 1E**), and the area of the lumen followed a similar pattern (**Fig. 1F**). The endometrium is also thickest at proestrus and thinnest at diestrus (**Fig. 1G**). However, the myometrial area is not significantly different across the estrous cycle (**Fig. 1H**). In addition to the area of these compartments, we also quantified the number of uterine glands and the luminal epithelial cell height across the estrous cycle. The number of glands is highest at proestrus and lowest at diestrus (**Fig. 1I**). Similarly, the luminal epithelial cell height is highest at proestrus and lowest at diestrus (**Fig. 1J**). Therefore, the majority of uterine compartments undergo remodeling across the estrous cycle. While the myometrial area does not remodel, the myometrium displays cyclical spontaneous contractile activity during the estrous cycle ^40^, so molecular or cellular remodeling may occur.

**Figure 1.**
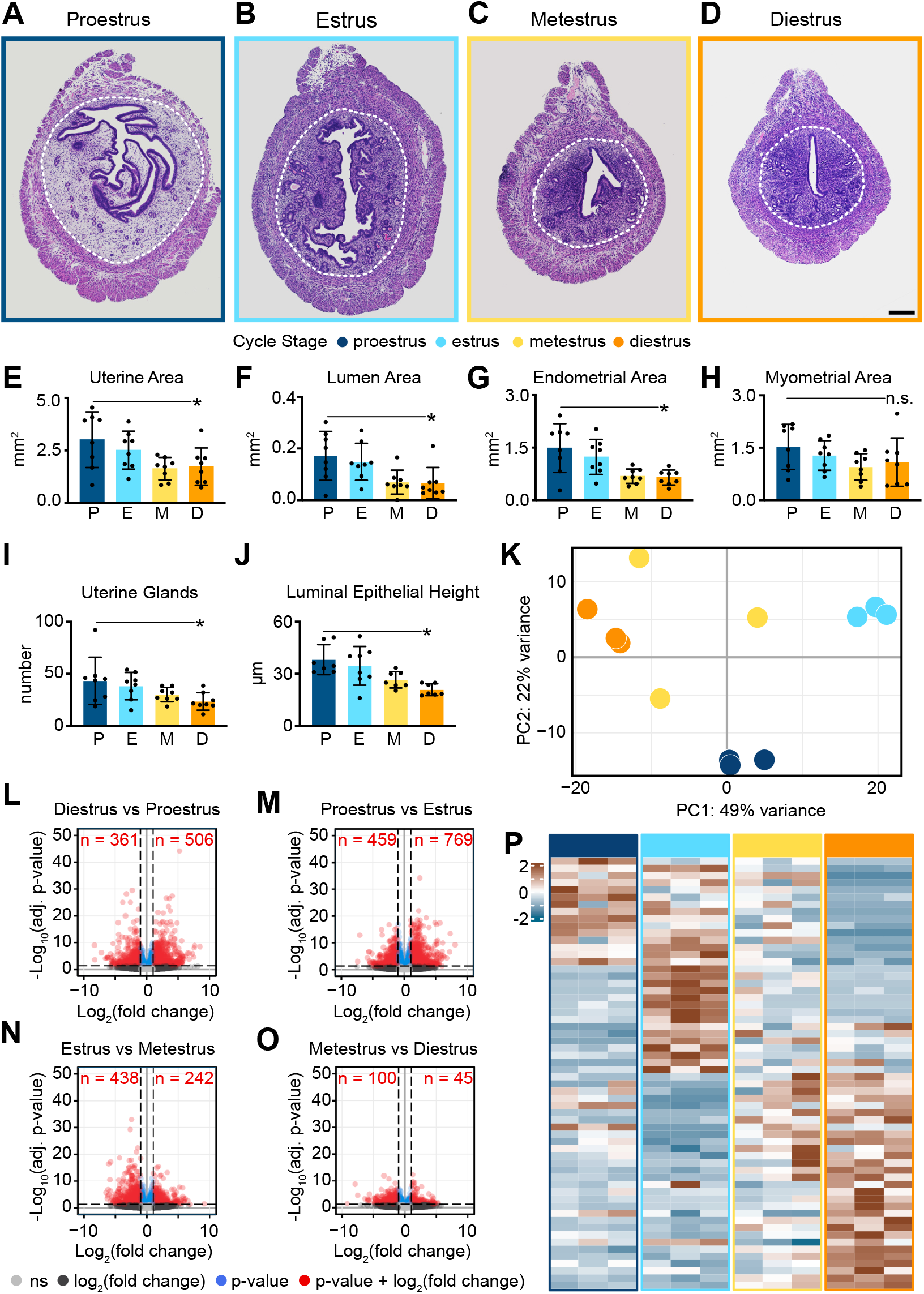
Cyclical mouse uterine remodeling occurs concomitant with transcriptional changes. H&E staining of mouse uterine sections at A) proestrus, B) estrus, C) metestrus, and D) diestrus where the dotted white line denotes boundary between the endometrium and myometrium. Quantitation of E) uterine area, F) lumen area, G) endometrial area, H) myometrial area, I) uterine gland number, and J) luminal epithelial height during those same timepoints. K) Principal Component Analysis (PCA) plot of 3’ TagSeq dataset. Volcano plots showing differential gene expression between L) diestrus and proestrus, M) proestrus and estrus, N) estrus and metestrus, and O) metestrus and diestrus. P) Heatmap of the most variable differentially expressed genes at each estrous cycle stage. All heatmaps use z-scores of the DESeq2 variance stabilized counts. * = p<0.05; n.s. = not significant.

### The murine uterus is transcriptionally dynamic across the estrous cycle

We hypothesized that downstream of steroid hormones, developmental signaling pathways would be re-deployed in the adult uterus to regulate cyclical uterine remodeling. To test this hypothesis, we performed 3’ TagSeq of the whole uterus at each stage of the estrous cycle (n=3 per stage). Principle component analysis (PCA) of three biological replicates per estrous cycle stage showed clear separation of estrous cycle replicates, with PC1 and PC2 accounting for 49% and 22% of the variance, respectively (**Fig. 1K**). While most samples within each stage cluster well together, we noted that the metestrus samples (in yellow) do not, which we expect is due to either biological variability or inaccurate estrous stage prediction.

Regardless, the stages clearly form a cycle within the PCA plot itself, with proestrus at the center bottom, estrus towards the middle right, metestrus towards the center top, and diestrus towards the middle left (**Fig. 1K**).

To take a more granular view of transcriptional changes during the estrous cycle, we specifically looked for differentially expressed genes at cycle transitions – for example, from diestrus to proestrus; proestrus to estrus; estrus to metestrus; and metestrus to diestrus. At most cycle transitions, there are hundreds of differentially expressed genes. At the diestrus-proestrus transition, 361 genes are downregulated and 506 are upregulated (**Fig. 1L**). At the proestrus-estrus transition, 459 genes are downregulated and 769 genes are upregulated (**Fig. 1M**). At the estrus-metestrus transition, 438 genes are downregulated and 242 genes are upregulated (**Fig. 1N**). Finally, at the metestrus-diestrus transition, 100 genes are downregulated and 45 genes are upregulated (**Fig. 1O**). To better visualize a subset of these differentially expressed genes (DEGs), we plotted the top 15 enriched DEGs of each cycle stage in a heatmap showing a cascade of gene expression during the estrous cycle (**Fig. 1P**). Therefore, the transcriptional landscape of the uterus is dynamically changing during the estrous cycle.

### Genes associated with embryonic development are re-deployed in the adult mouse uterus during the estrous cycle

We were curious if the DEGs at each cycle transition were enriched in particular gene classes, so we performed Gene Ontology (GO) analysis ^41^. In support of our overarching hypothesis, developmental biology GO terms were prevalent at metestrus. The top GO terms included: limb development; appendage development; limb morphogenesis; appendage morphogenesis; tube development; and embryonic limb morphogenesis (**Fig. 2C**). At proestrus, the top GO terms included sensory perception of smell; sterol biosynthetic process; and epithelium development (**Fig. 2A**). At estrus, the top GO terms included response to external stimulus; and response to other organism (**Fig. 2B**).

**Figure 2.**
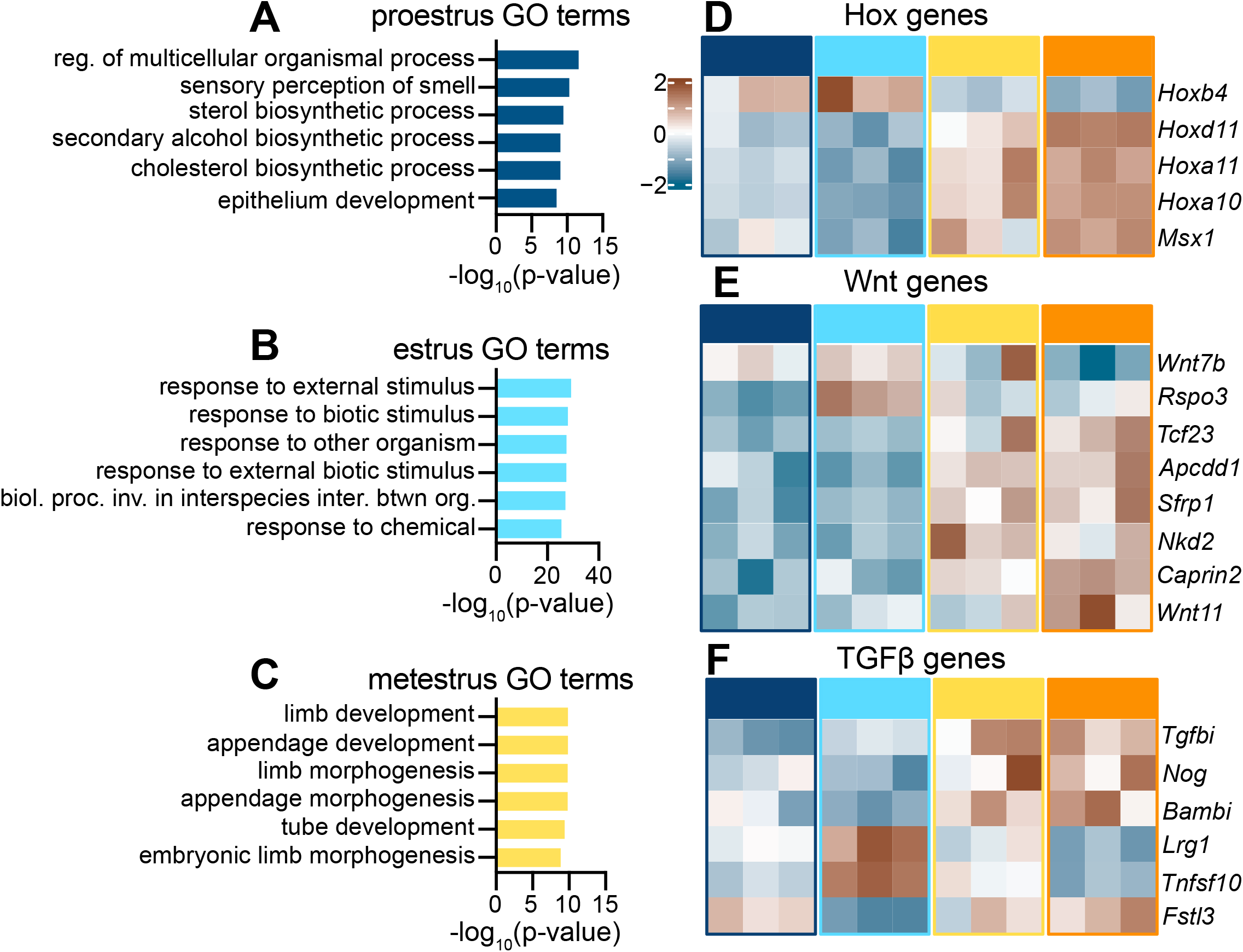
Development genes are cyclically expressed in the uterus during the estrous cycle. GO terms of differentially expressed genes enriched at A) proestrus, B) estrus, and C) metestrus. Heatmaps of D) Hox genes, E) Wnt signaling, and F) TGFβ signaling. All heatmaps use z-scores of the DESeq2 variance stabilized counts.

Using the GO term analysis as a partial guide, we manually curated the developmental genes and signaling pathways that were differentially expressed in the uterus across the estrous cycle. For example, Hox genes and *Msx1* are required for implantation, but their role during the estrous cycle is unknown ^42-45^, and we found that homeobox genes were differentially expressed: *Hoxa10* and *Hoxa11* are upregulated when progesterone is high during metestrus and diestrus, along with *Hoxd11* and *Msx1* (**Fig. 2D**). In contrast, *Hoxb4* is upregulated at proestrus and estrus when estrogen is high (**Fig. 2D**). Likewise, *Wnt7b* is required for uterine gland development ^46,47^, and dynamic spatial expression of other Wnts has been reported during the estrous cycle ^48^. We found that *Wnt7b* and *Wnt11* are differentially expressed across the estrous cycle (**Fig. 2E**), and *Rspo3* is upregulated specifically at estrus, while other Wnt signaling genes, *Apcdd1, Sfrp1, Nkd2*, and *Caprin2* are enriched at metestrus and diestrus (**Fig. 2E**). In addition to Hox and Wnt signaling genes, TGFβ signaling is also required for implantation and uterine gland and myometrial development ^49,50^, but it’s role during the estrous cycle is unknown. In our dataset, we also identified differential expression of genes involved in TGFβ signaling. For example, *Tgfbi, Nog*, and *Bambi* are enriched at metestrus and diestrus, *Lrg1* and *Tnfsf10* are enriched at estrus, and *Fstl3* is enriched at both diestrus and proestrus (**Fig. 2F**). Thus, the expression of many of genes required for either implantation or uterine development, are dynamically expressed across the estrous cycle, though the relationship between this cyclical expression and cyclical uterine remodeling has not been discovered.

### Hedgehog signaling components are cyclically co-regulated across the estrous cycle

Our most striking finding was that the genes encoding nearly all members of Hedgehog signaling pathway (**Fig. 3A**) were cyclically expressed together and enriched at diestrus (**Fig. 3B-M**). The major Hh ligand in the uterus, *Ihh*, and the known downstream target genes of the pathway, such as *Gli1* and *Ptch1*, are significantly enriched (denoted with an asterisk) (**Fig. 3B**). Given that *Ihh* is a direct target of progesterone at implantation ^6,8,51^, enrichment of *Ihh* at metestrus or diestrus when progesterone is high was expected ^3^. However, the remaining signaling components, including *Gli2, Gli3, Ptch2, Smo, Sufu, Gpr161, Kif7, Evc* and *Evc2* are not typical downstream targets of Hh signaling, yet they also trend upward at diestrus (**Fig. 3B**). We confirmed the significant enrichment of the majority of these genes by qPCR on an additional 5 animals for a total of n=8 (**Fig. 3C-M**). Thus, uterine expression of the genes encoding most of the Hh signaling cassette are co-regulated across the mouse estrous cycle.

**Figure 3.**
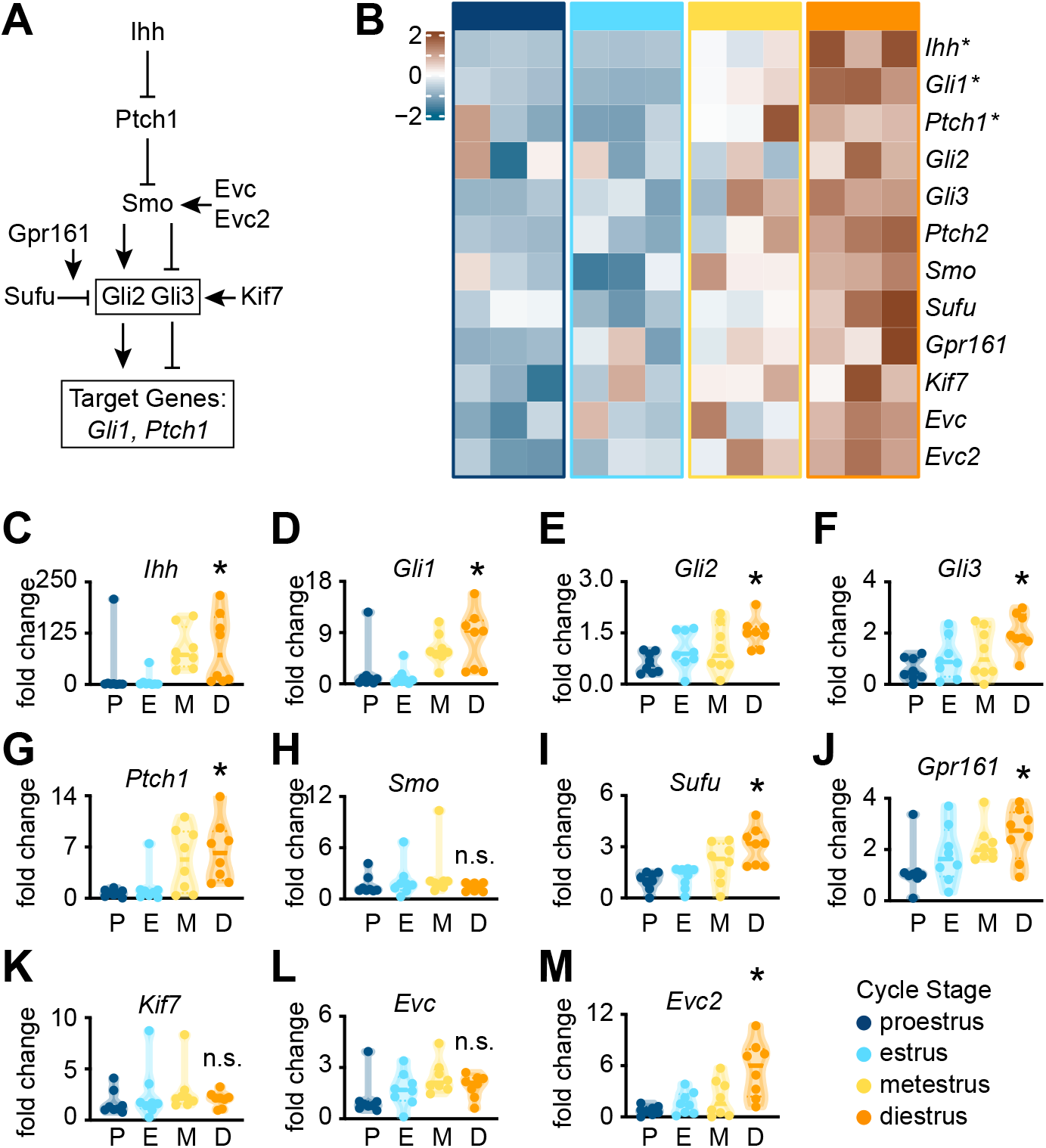
The Hedgehog signaling pathway is cyclically regulated across the estrous cycle. A) The biochemical pathway of Hh signaling. B) Heatmap of the core components of Hh signaling across the estrous cycle, where the z-score of the DESeq2 variance stabilized counts is used. CM) qPCR of the same core components of Hh signaling with n=8 samples. * = p<0.05; n.s. = not significant.

### Hedgehog signaling is required for iterative uterine remodeling across the estrous cycle

While Hedgehog signaling has been extensively studied at implantation, specifically in the endometrium, nothing is known about possible roles for Hedgehog signaling in uterine remodeling or homeostasis across the estrous cycle. We therefore made a conditional knockout of the major activator, *smoothened* (*Smo*) using the *progesterone receptor cre* (*PR-Cre*, **Fig. 4A**) which is expressed throughout the adult uterus ^24,52^. The controls (ctrl) were either *PR-Cre*^*+/- or -/-*^ *Smo*^*fl/+ or +/+*^ and the conditional knockouts (*Smo* cKO) were *PR-Cre*^*+/-*^ *Smo*^*fl/fl*^. We confirmed that *Smo* is indeed significantly decreased or absent in the cKOs compared to the controls by qPCR (**Fig. 4B**). *Smo* is the major activator of the Hh signaling pathway, therefore, the *Smo* cKOs should not be able to activate downstream Hh signaling target genes. We confirmed this by checking for the expression of *Gli1*, a downstream target gene of the pathway. Indeed, while *Gli1* is upregulated at diestrus compared to estrus in our controls, that fails to occur in the cKOs (**Fig. 4C**). In addition to looking at the expression of *Smo* and *Gli1*, we also looked at fertility because previous work has shown that Hedgehog signaling is required for implantation ^8,10,11,51,53^. To confirm that our cKO is infertile, we performed breeding studies on the controls and *Smo* cKOs. The control mice had an average of 9 pups per litter, while the *Smo* cKOs failed to ever have pups (**Fig. 4D**). Finally, we asked whether the *Smo* cKOs are cycling through the estrous cycle using vaginal cytology. We show that the cKOs are able to progress through the cycle, albeit with a significantly shorter diestrus stage (**Fig. 4E**).

**Figure 4.**
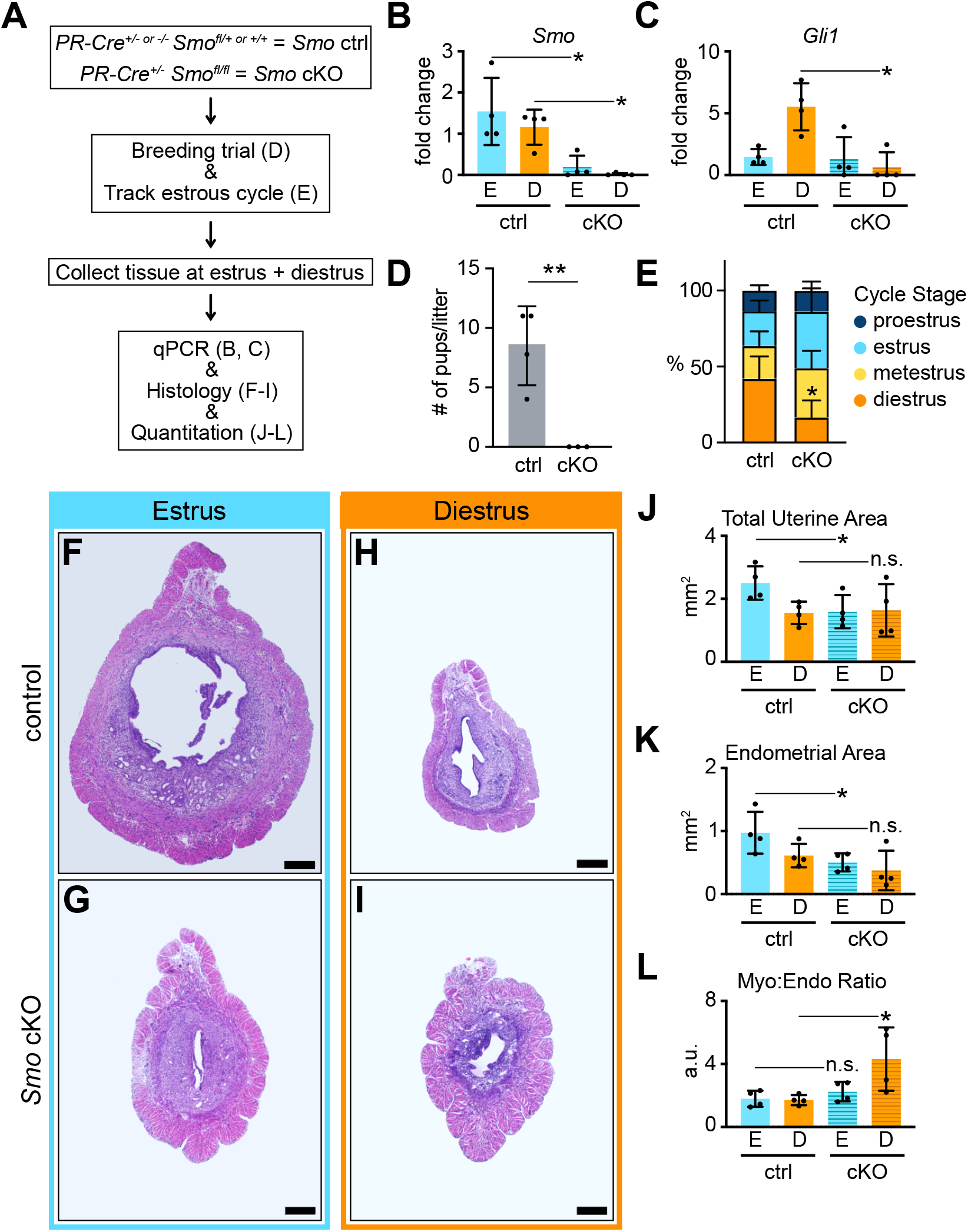
Hedgehog signaling is required for cyclical uterine remodeling. A) Experimental flow to determine role of Hh signaling in the adult uterus during the estrous cycle. B) qPCR for *Smoothened* in the controls and *Smo* cKOs. C) qPCR for *Gli1* in the controls and *Smo* cKOs. D) Breeding trial of controls and *Smo* cKOs. E) Estrous cyclicity of controls and *Smo* cKOs. F-I) H&E stained uterine sections from control and *Smo* cKOs. Quantitation of J) total uterine area, K) endometrial area, and L) myometrial-to-endometrial area ratio. * = p<0.05; n.s. = not significant.

Our control experiments showed that our *Smo* cKO behaved as we would expect, so we next investigated its effect on cyclical uterine remodeling. We focused on two stages: estrus (E) is in the first half of the cycle when estrogen is high, whereas diestrus (D) is the last stage of the cycle when progesterone is enriched ^3^. In the controls, the uterus is larger at estrus compared to diestrus (**Fig. 4F** compared to **Fig. 4H**, quantified in **Fig. 4J**). In the *Smo* cKO at estrus, the uterus is significantly thinner compared to controls (**Fig. 4G** compared to **Fig. 4F**, quantified in **Fig. 4J**). This change in overall uterine size at estrus is likely due to a decrease in the endometrial area (**Fig. 4K**). At diestrus, the *Smo* cKO is still thin (**Fig. 4I, J**). And while the endometrium is still thin (**Fig. 4K**), we observed that the ratio of myometrium to endometrium was different. We quantified the ratio of myometrial area to endometrial area, and in the *Smo* cKO at diestrus, that ratio is significantly higher compared to the other samples (**Fig. 4L**). Thus, Hedgehog signaling is required for normal cyclical remodeling of the endometrium and homeostasis of the myometrium across the estrous cycle.

### Hedgehog signaling maintains the typical morphology of the longitudinal smooth muscle layer

Our data suggest that one of the key sites of action of Hh in the cycling uterus was the myometrium, so we more closely observed the morphology, including thickness and folding of the *Smo* cKO myometrium. There are two layers of smooth muscle in the uterus: the inner circular layer and the outer longitudinal layer. In the *Smo* cKO at diestrus it was the longitudinal layer that appeared thicker so we quantified it by drawing a straight line from the outer edge of the circular muscle to the outer edge of the longitudinal muscle. At diestrus, the *Smo* cKO longitudinal smooth muscle trends towards increased thickness compared to the other samples, although isn’t statistically significant (**Fig. 5A**). We reasoned that the longitudinal layer may look thicker because it is more folded compared to controls (**Fig. 5B-E**). We therefore calculated ‘smoothness’ as a proxy for folding, defined by the ratio of the uterine perimeter length (**Fig. 5B’**) to the best fit ellipse (**Fig. 5B’’**). While there is no significant difference between the controls at estrus or diestrus, there is a trend towards a decrease at estrus between the control and *Smo* cKO (**Fig. 5B** compared to **Fig. 5D**, quantified in **Fig. 5F**), and a significant decrease at diestrus between the control and *Smo* cKO (**Fig. 5C** compared to **Fig. 5E**, quantified in **Fig. 5F**). These data suggest that the *Smo* cKO myometrium, specifically at diestrus, is less smooth and therefore atypically folded compared to controls.

**Figure 5.**
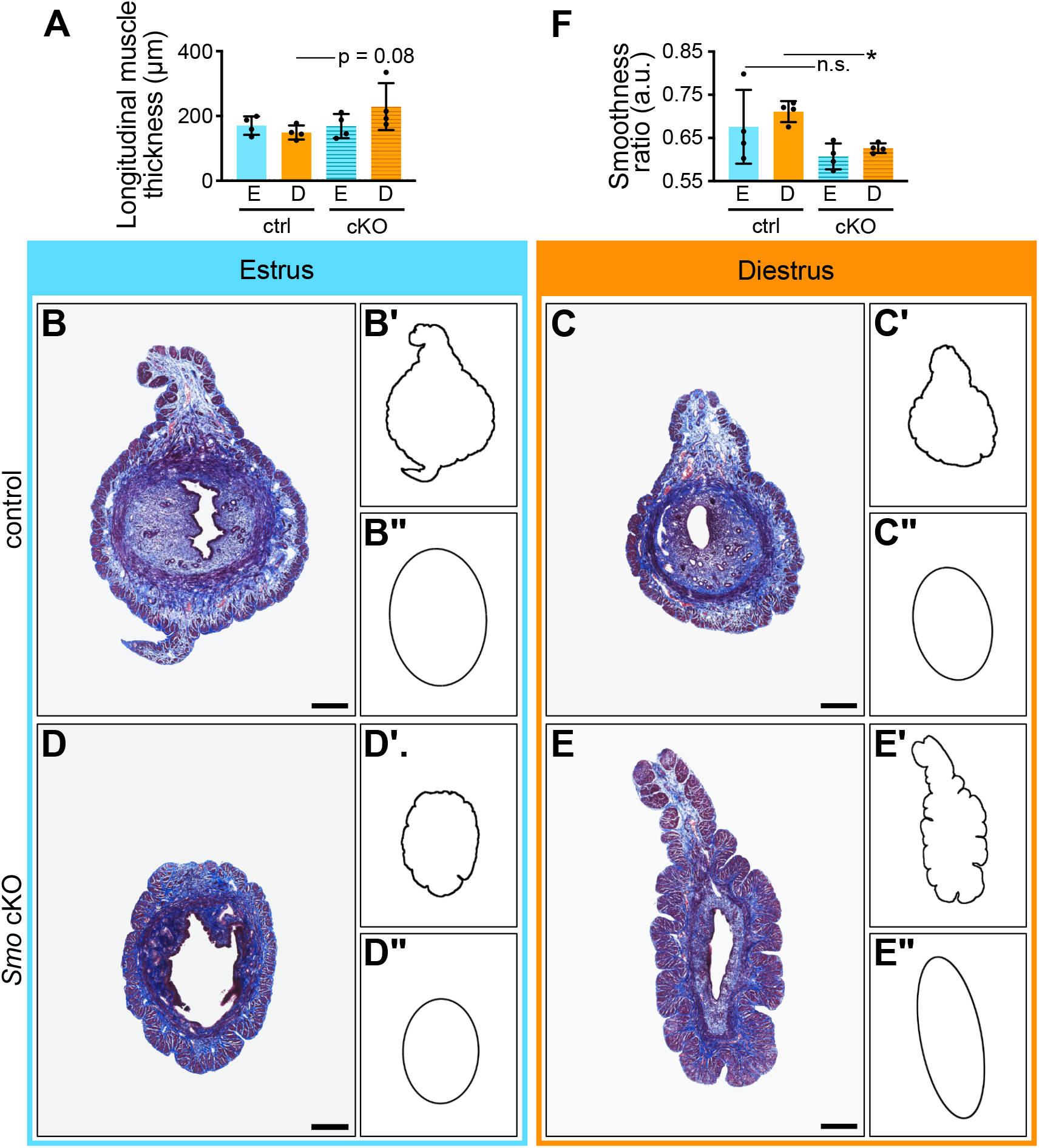
The myometrium is significantly folded in the absence of Smoothened. A) Quantitation of longitudinal muscle thickness. B-E) Masson Trichrome histology of the mouse uterus. B’-E’) Outline of sections highlighting the ruffled nature of the *Smo* cKO. B”-E”) Best fit ellipse of the outline. F) The smoothness of each sample was determined by taking the ratio of the true outline to the best fit ellipse. * = p<0.05; n.s. = not significant.

### Muscle fibers are larger in the absence of Smoothened

Folding similar to what we see in the myometrium of *Smo* cKO at diestrus has been reported to occur naturally in some developing organs. For example, in some mammalian brains, the exterior of the brain becomes highly folded and there is evidence that this results from the outer grey matter growing more than the inner white matter, resulting in mechanical buckling ^54^. The inverse is true in the developing vertebrate gut, where the growing inner endoderm and mesenchyme are restrained by the outer layers of smooth muscle resulting in folding ^55^. We hypothesized that the folding in the *Smo* cKO at diestrus may be caused by an increase in longitudinal muscle fiber number or size compared to the unchanging endometrial area (**Fig. 4K**).

To quantify the size and number of muscle fibers, we used Masson Trichrome histology because it stains muscle in red, and collagen in the extracellular matrix in blue (histology in **Fig. 5** and **Fig. 6**). Given that uterine smooth muscle fibers are embedded in extracellular matrix, ^56^, we were able to assess smooth muscle fibers using Ilastik which takes advantage of color and texture differences to differentiate structures ^27^. Here, Ilastik produced masks of muscle fibers (**Fig. 6A-D, Fig. 6A’’-D’’**). The average fiber number was similar in each of our conditions (**Fig. 6E**). The average fiber size was unchanged between the controls, and significantly increased in the *Smo* cKO at diestrus (**Fig. 6F**). We binned fibers by area and found that the smallest fibers were significantly decreased in the *Smo* cKO at diestrus, with a concomitant significant increase in the largest fiber size (>6,001μm^2^, **Fig. 6G**). Therefore, one mechanism by which smooth muscle folding may occur is an increase in fiber size in the *Smo* cKO uteri at diestrus.

**Figure 6.**
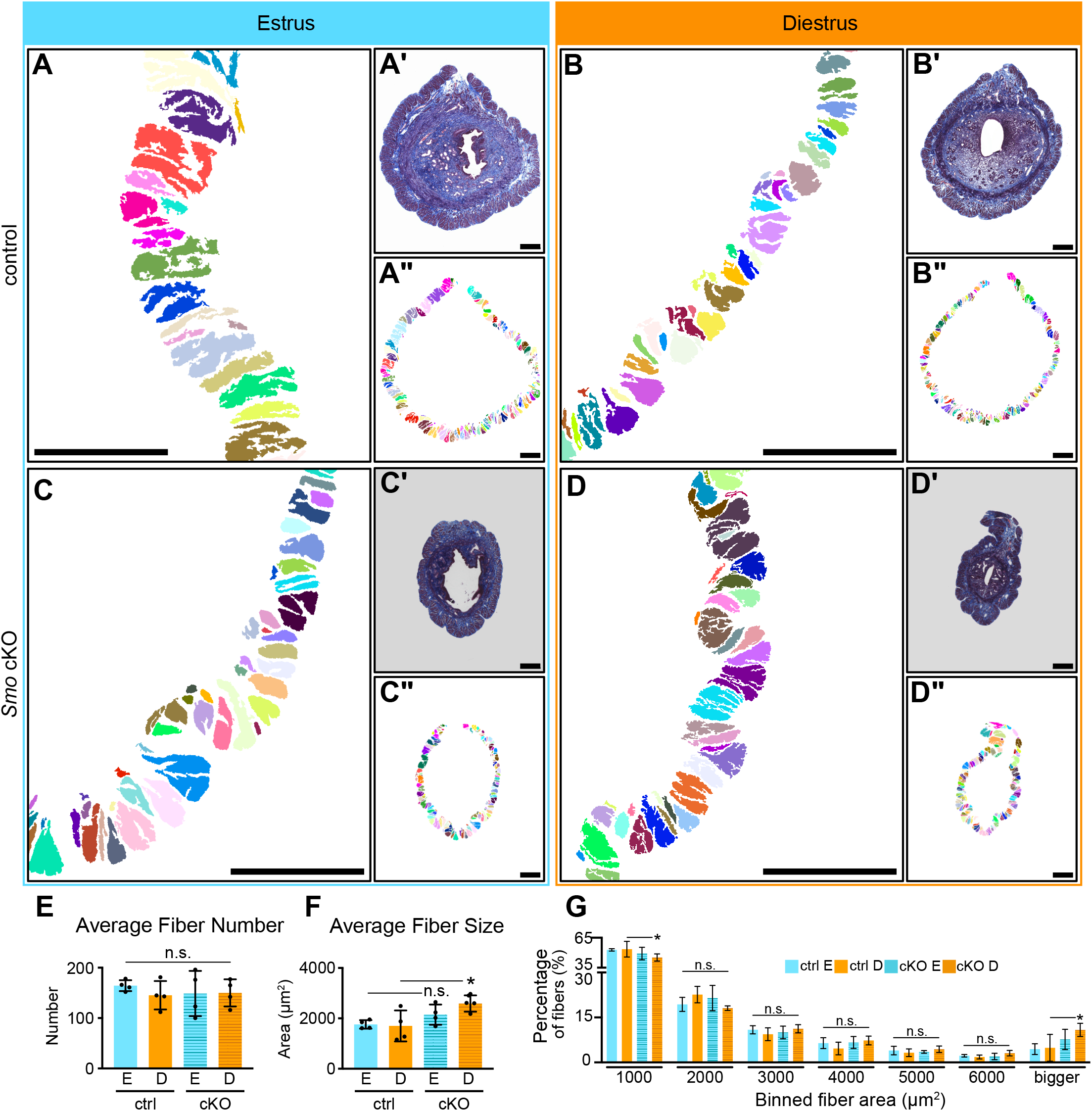
Myometrial fibers are significantly bigger in the absence of Smoothened. A-D) Masks of the longitudinal muscle fibers were produced using Ilastik. A’-D’) Masson Trichrome histology was used to identify the myometrial muscle fibers. A”-D”) Longitudinal fibers were identified throughout the entire section. E) Quantitation of the fiber number in each sample. F) Quantitation of the fiber area in each sample. G) Fiber areas were binned in 1000μm^2^ increments. * = p<0.05; n.s. = not significant.

## Discussion

One of the first studies to show that the human uterus undergoes typical cyclical remodeling during the menstrual cycle rather than pathological changes, was published in 1908 ^57,58^. Since then, a large body of literature has characterized the regeneration that occurs, and has focused on understanding menstruation as well as timing of re-epithelialization of the uterus ^33,59-63^. However, it would be nearly 100 years before a similar study on mouse uterine remodeling relative to circulating steroid hormone levels was published ^3^, despite this animal’s power and widespread use as a tractable genetic model for mammalian biology.

In addition, mouse models have been used extensively to understand the impact of estrogen and progesterone on the uterus, but almost exclusively by exogenously administered hormones to ovariectomized mice^64,65^, genetically modified mice to examine loss of steroid hormone signaling ^5,66-68^, or at implantation ^8,12,69-72^. In contrast, far less molecular work has been done to understand the normally cycling mouse uterus. Such work is important because it provides a framework to begin assessing regeneration and remodeling in other systems and the mouse is the only major genetically tractable mammalian model system.

To start developing the mouse as a model for cyclical uterine remodeling, we determined the cyclical transcriptome of the whole uterus and found that hundreds of genes are differentially expressed at each cycle transition. Given that each estrous cycle stage is on average 24 hours long, our data suggest that the mouse uterus is undergoing radical transcriptional changes within that time frame. This is contrast to the mouse oviduct, which we previously showed undergoes minimal transcriptional change across the estrous cycle ^73^.

We initially hypothesized that aspects of cyclical uterine remodeling would be driven by re-deployment of developmental signaling pathways and genes. Gratifyingly, many developmental pathways are differentially expressed across the estrous cycle in the mouse uterus, including Hox genes, Wnt signaling, and TGFβ signaling. Components of these signaling pathways are differentially expressed in the mouse or human uterus across the estrous or menstrual cycle ^48,74-77^. All three are also required for uterine development and implantation ^46,78-84^. Wnt and TGFβ signaling all play a role in uterine gland development ^49,85^, so it is possible that they control various aspects of uterine gland morphogenesis across the estrous cycle.

Interestingly, dysfunctional TGFβ signaling results in excess smooth muscle specification during development ^49,86^, so it may also play a role in myometrium homeostasis during the estrous cycle. However, a functional role for these developmental signaling pathways and genes on the cycling uterus is unknown.

We focused on Hedgehog (Hh) signaling because most Hedgehog signaling components were co-regulated and elevated together at diestrus, and it is a fundamental pathway that regulates embryonic development and adult tissue homeostasis. Hh signaling is required for uterine development as well as endometrial function at implantation ^8,10,51,53,70,87-89^. In other organs, Hh signaling plays a role in smooth muscle differentiation during development and homeostasis ^89-95^. Finally, in adults, Hh signaling is instrumental in controlling remodeling or regeneration of hair follicles and skeletal muscle ^29,88,96^. The adult uterine endometrium remodels and the myometrium is homeostatic across the estrous cycle, yet nothing is known about Hh signaling in these uterine compartments during this timeframe.

To fill these knowledge gaps, we conditionally deleted *smoothened* using the *progesterone receptor Cre*. Loss of *Smo* in the cycling uterus resulted in a lack of endometrial remodeling across the estrous cycle. At implantation, loss of *Ihh* results in lack of stromal cell proliferation and therefore implantation failure ^8^. The role of Hh signaling in the endometrium across the estrous cycle may be similar to what has been previously published at implantation, but additional follow up experiments will be necessary to confirm. In addition to the endometrial phenotype, we also documented a myometrial phenotype: in the absence of *Smo* the myometrium has increased folding, which may be caused by the significant increase in smooth muscle fiber size at diestrus. Whether this increase in fiber size is due to hyperplasia or hypertrophy remains to be determined.

Our analysis focused on mice that are cycling, not pregnant. However, throughout gestation, myometrial muscle fibers undergo hyperplasia followed by hypertrophy to account for fetal growth and prepare for labor ^97^. During involution, or the healing that occurs post-birth, the myometrium undergoes remodeling to return to a non-pregnant state. In an H&E stained postpartum day 1 mouse uterine section, the myometrium looks folded similar to the *Smo* cKO at diestrus ^98^. Our data suggest Hh signaling may be important for myometrial remodeling that occurs during late pregnancy or involution, and warrants a robust study into myometrial folding during pregnancy, parturition, and involution.

Finally, our study has broader implications for human health and disease. Hh signaling is upregulated in a variety of uterine pathologies like uterine smooth muscle tumors ^99-101^, and endometriosis ^102-104^. However, whether Hh signaling plays a role in initiation or progression of these pathologies remains unclear.

## Acknowledgements

The authors are deeply grateful to the staff of the Genomic Sequencing and Analysis Facility and the Animal Resource Facility at the University of Texas at Austin. We would also like to thank Dr. Carmen Williams, Dr. Stephen Vokes, and the members of the Wallingford lab for critical scientific discourse relevant to the data presented here. This work was supported by the NIH/NICHD: F32HD095618 (E.C.R.) and R01HL117164 and R01HD085901 (J.B.W.). This work was also supported by the University of Texas at Austin Undergraduate Research Fellowships (K.T. and H.M.).

## Author Contributions

**E.C.R**.: Conceptualization, Methodology, Investigation, Data curation, Formal Analysis, Visualization, Writing, Supervision, Project Administration. **N.K.T**.: Investigation, Validation. **A.N.G**.: Investigation, Validation. **H.M**.: Investigation, Validation. **M.N**.: Investigation, Validation. **T.R**.: Investigation, Validation. **R.D.F**.: Investigation. **J.B.W**.: Conceptualization, Writing – Review & Editing, Supervision, Funding Acquisition, Project Administration.

## Supplemental Table

**Table S1.**
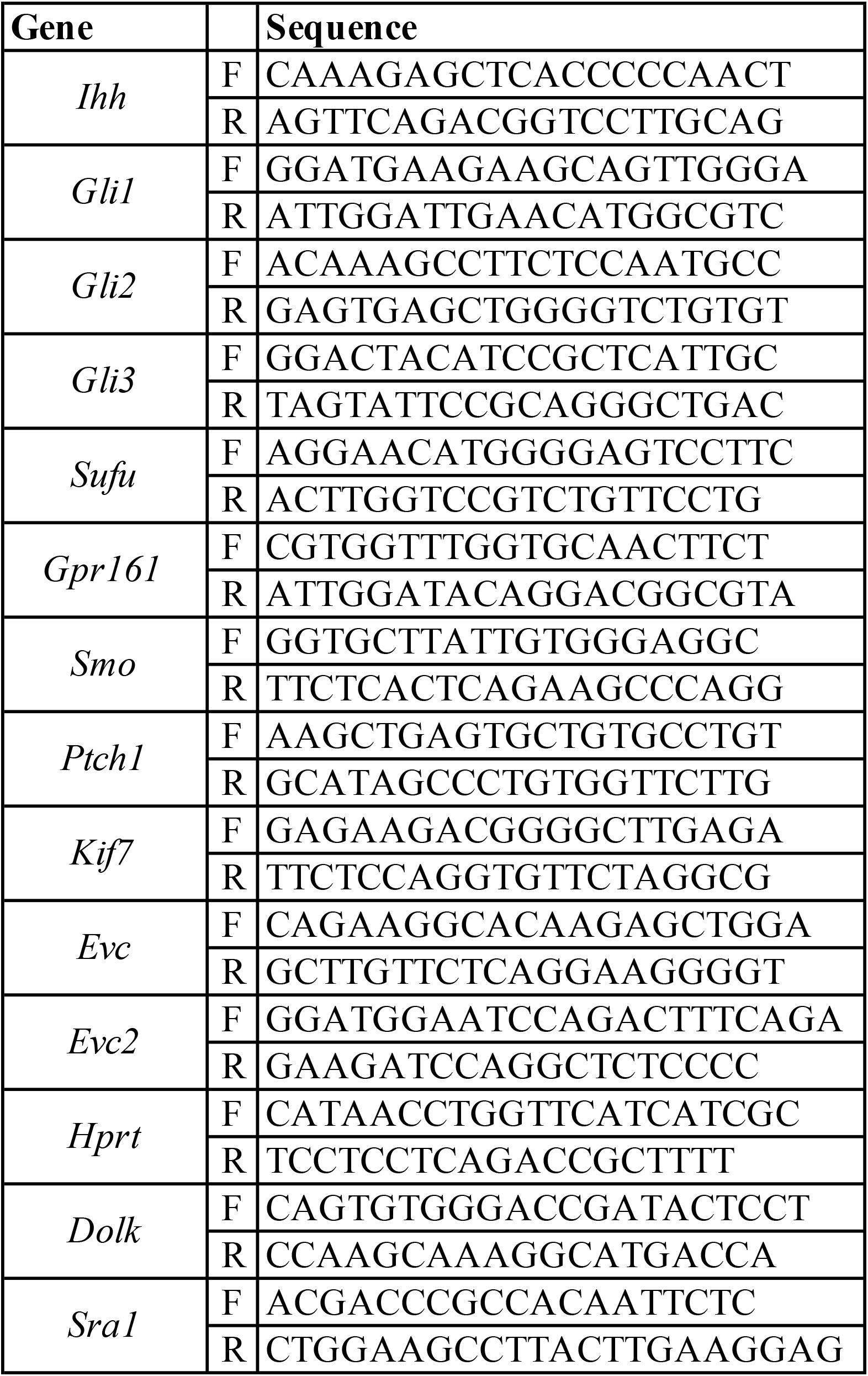
Primers used for qPCR.

